# Heritability of hippocampal functional and microstructural organisation

**DOI:** 10.1101/2021.11.10.468049

**Authors:** Şeyma Bayrak, Reinder Vos de Wael, H. Lina Schaare, Meike D. Hettwer, Benoit Caldairou, Andrea Bernasconi, Neda Bernasconi, Boris C. Bernhardt, Sofie L. Valk

## Abstract

The hippocampus is a uniquely infolded allocortical structure in the medial temporal lobe that consists of the microstructurally and functionally distinct subregions: subiculum, cornu ammonis, and dentate gyrus. The hippocampus is a remarkably plastic region that is implicated in learning and memory. At the same time it has been shown that hippocampal subregion volumes are heritable, and that genetic expression varies along a posterior to anterior axis. Here, we studied how a heritable, stable, hippocampal organisation may support its flexible function in healthy adults. Leveraging the twin set-up of the Human Connectome Project with multimodal neuroimaging, we observed that the functional connectivity between hippocampus and cortex was heritable and that microstructure of the hippocampus genetically correlated with cortical microstructure. Moreover, both functional and microstructural organisation could be consistently captured by anterior-to-posterior and medial-to-lateral axes across individuals. However, heritability of functional, relative to microstructural, organisation was found reduced, suggesting individual variation in functional organisation of subfields is under low genetic control. Last, we demonstrate that structure and function couple along its genetic axes, suggesting an interplay of stability and plasticity within the hippocampus. Our study provides new insights on the heritability of the hippocampal formation and illustrates how genetic axes may scaffold hippocampal structure and function.

## Introduction

The hippocampal formation in the medial temporal lobe is involved in numerous functions such as episodic memory ^1–3^, spatial navigation ^4^, emotional reactivity ^5^ and stress resilience ^6–8^. It is a region highly susceptible to disorder in various neurological and neuropsychiatric conditions, such as schizophrenia ^9^, posttraumatic stress disorder ^10^, temporal lobe epilepsy ^11^, and Alzheimer’s disease ^12^. Having a three layered allocortex, the hippocampal formation consists of multiple subfields, or zones, starting at the subiculum (SUB) and moving inward to the hippocampus proper; the cornu ammonis (CA), and dentate gyrus (DG) ^13–16^. These subfields have unique microstructure ^15,16^ and participate differently in the hippocampal circuitry ^17^, likely implicating different contributions to function ^18–20^. Beyond the internal hippocampal wiring, anatomical projections to isocortical targets vary based on the position within the hippocampal formation ^21,22^. Thus, the intrinsic organisation of the hippocampus relates to its connectivity to the rest of the brain. For example, tracer studies in rodents have shown that the ventral hippocampus is anatomically connected to the olfactory regions, prefrontal cortex, and amygdala, while the dorsal hippocampus is connected to the retrosplenial cortex, mammillary bodies, and anterior thalamus ^23,24^. This ventral-dorsal transition in rodents may relate to an anterior-posterior (A-P) axis in humans ^25,26^. Conversely, hippocampal infolding aligns with a medial-lateral (M-L) axis followed by the subfields, suggesting another transitional axis driven by intracortical microstructure ^16,27^. Thus, the hippocampal formation features two major axes, one from anterior to posterior segments, and the other along its infolding from SUB via CA to DG.

Hippocampal organisational axes can be described using gradients ^28^. This framework enables continuous representations of the high-dimensional inter-regional patterns, unrestricted by the traditional network boundaries ^29^ 30. Along each gradient axis, voxels/vertices sharing similar connectivity patterns are situated close to each other, whereas those most divergent are at opposite ends of the respective axis ^31^. Using this method, hippocampal organisational axes observed in the structure of the hippocampus have been reported to be paralleled by the functional organisation of the hippocampus, as measured *in vivo* using functional MRI ^32 33–35^. Hippocampal gradients were further associated with its microstructural organisation ^35^, as well as performance on memory recollection ^34^ and pattern separation tasks ^33^, suggesting a link between functional organisation of the hippocampus, its structure, and behavioural variability. At the same time, whether hippocampal functional and microstructural organisation axes vary according to genetic factors or rather adapt flexibly as a function of environment is incompletely understood.

There is ample evidence for both genetic and environmental factors playing a role in hippocampal organisation. For example, in parallel to observations in rodents, it has been shown that the human hippocampus shows variations in gene expression along the A-P axis ^36^. This transcriptomic gradient is related to different macroscale networks in the cortex, dissociating behavioural systems associated with action and spatial cognition on the one hand and social and emotion/motivation on the other. Paralleling observations of differential gene expression, recent work has indicated that hippocampal subfields have differentiable genetic signatures ^37^ providing further evidence for the complex molecular biology associated with the hippocampal formation. Hippocampal subfield volumes have also been shown to be heritable ^38^, indicating individual variations of subfield volume are, in part, under genetic control. However, whether the internal structural and functional organisation of hippocampal subfields is heritable, or rather varies as a function of non-genetic factors, is not known to date. Indeed, other lines of research have reported the hippocampus to be highly plastic and reactive to stress ^39,40^ 41. One way to reconcile the notion of plasticity and stability reported in the hippocampus is by means of the structural model ^42^. This model links isocortical cytoarchitecture and associated connectivity patterns to regional variations in plasticity and stability ^43–46^. In the isocortex, structure-function coupling has been shown to progressively decrease along an axis from unimodal to transmodal regions ^43–46^. Such uncoupling is paralleled by reductions in genetic control from unimodal to transmodal regions ^43^ and may facilitate more flexible computations enabling abstract human cognition ^47,48^. It is possible that the coupling of microstructure and function in the hippocampus shows meaningful variation along its large-scale axes, and helps to further understand the interrelation between plasticity and stability, or genetic and environmental factors, within this allocortical structure.

Here, we studied how the genetically controlled organisation of the hippocampus may support learning and plasticity. To do so, we leveraged the multimodal dataset of the Human Connectome Project ^49^ to sample resting state functional time series as well as T1w/T2w intensity (as a proxy for intracortical myelination ^50^) in both the hippocampal subfields and isocortex. Diffusion map embedding ^51^ was used to describe the largest axes of variance in functional connectivity and microstructural covariance ^25,35^. The twin set-up of the HCP data enabled us to quantify both the heritability of these intrinsic functional and microstructural representations, as well as the genetic coupling between hippocampal and isocortical microstructural profiles. Last, we studied the shared organisation of hippocampal function and structure to describe spatial co-variation of structure-function associations along genetic hippocampal organisational axes. We performed extensive robustness analysis to assess the stability of our findings. Overall, these analyses will help to further understand the relationship between the genetic basis of hippocampal organisation and its flexible functional role.

## Results

### Hippocampal-isocortical functional connectivity is heritable (Figure 1)

Hippocampal subfields i.e. subiculum (SUB), CA1-3 (CA), and CA4-DG (DG), were delineated automatically using SurfPatch, a previously validated surface-based subfield segmentation algorithm ^52^ (**Fig. 1A**). Resting-state (rs) fMRI time series were extracted along subfield surfaces and isocortical parcels (Glasser Atlas of 360 areas ^53^) and correlated to estimate functional connectivity (FC). After quality assessment (**Fig. S1A**), n = 709 participants (395 women, mean ± SD age = 28.7 ± 3.7 y, 176 monozygotic twins, 178 siblings without twin status and 355 participants without familial relatedness, HCP S900 data release ^49^) were included.

**Fig. 1.**
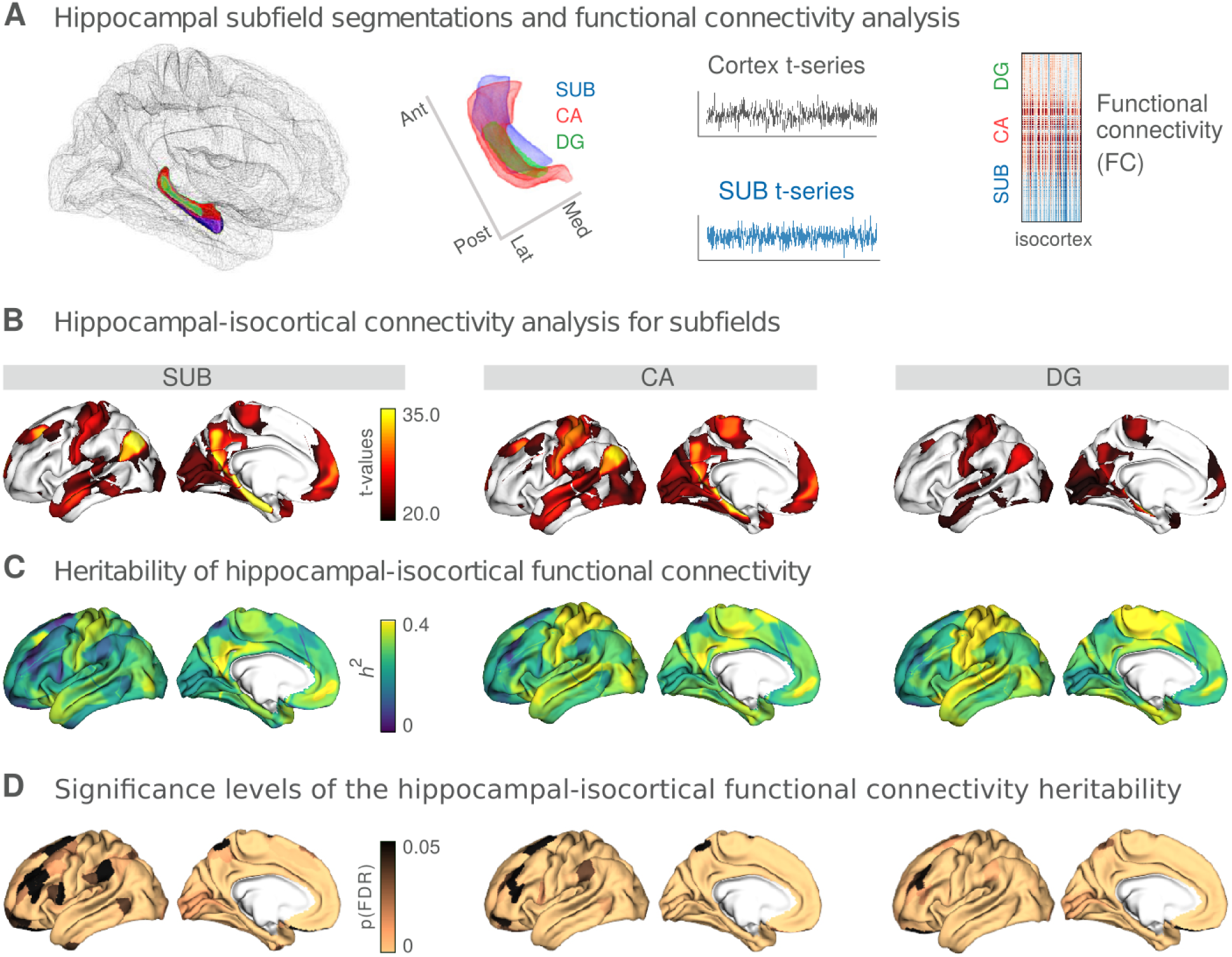
Hippocampal-isocortical functional connectivity and its heritability. **A**. Hippocampal subfield surfaces were automatically delineated using SurfPatch ^52^: subiculum (SUB, blue), CA1-3 (CA, red), and CA4-DG (DG, green). rs-fMRI time series were extracted along the individual subfields and correlated with the time series of the isocortex to obtain the functional connectivity (FC). **B**. Isocortex-wide FC of SUB (left), CA (middle), and DG (right). Isocortex-wide findings were thresholded at t > 20 to represent the highest connections. **C**. Heritability (*h*^2^) scores of the subfield-isocortical functional couplings throughout the cortex. **D**. Significance levels of the *h*^2^ scores from panel C. Significance level was reported with the multiple comparison corrected p-values (p(FDR)). Copper colour denotes pFDR < 0.05 and black colour pFDR ≥ 0.05.

Subfield-isocortical FC measures were mapped using linear and mixed effects models in BrainStat and thresholded at t > 20 to indicate highest connections (https://github.com/MICA-MNI/BrainStat ^54^) (**Fig. 1B**). The strongest connections were found in the default-mode, somatomotor, visual and limbic areas, across subfields. Heritability analyses (Sequential Oligogenic Linkage Analysis Routines, SOLAR, v8.5.1) ^55^ (*h*^2^) indicated that SUB-isocortex FC was the highest in regions part of sensorimotor (mean *h*^2^ score: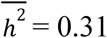) and default mode 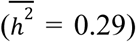 networks (**Fig. S1B**). A similar heritability profile was observed for CA-isocortex FC, with highest heritability in sensorimotor 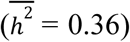, default mode 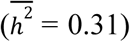 and dorsal attention 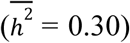 networks. For the DG-isocortex FC, compared to SUB and CA, we observed a higher heritability in the sensorimotor 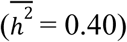 and ventral attention 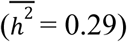 networks. The significance level of the *h*^2^ scores were assessed using a likelihood ratio test (p-values) and then corrected for multiple comparisons using FDR (pFDR) (**Fig. 1D**). Throughout most of the cortical parcels, the heritability was found to be significant, with an increasing number of significant parcels from SUB towards CA and DG.

### Hippocampal functional organisation is moderately heritable (Figure 2)

Following, we aimed to evaluate whether the functional organisation within hippocampal subfields was heritable as well. To do so, we first constructed topographic gradients of the hippocampal FC patterns using unsupervised dimension reduction ^35,51^ (**Fig. 2A**). Replicating previous work ^35^, the principal subfield gradient (*G*1_*FC*_) presented an A-P axis across hippocampal subfields and explained 24% of the variance, whereas the second subfield gradient (*G*2 _*FC*_) described a M-L axis and explained 9% of the variance (**Fig. S2A**). Anterior hippocampal subfield portions (blue in **Fig. 2A**) were functionally coupled to sensorimotor, default mode and limbic networks (**Fig. 2B, Fig. S2B**). Posterior hippocampal subfield portions (yellow in **Fig. 2A**) were functionally more connected to fronto-parietal, salience, dorsal attention and visual networks (**Fig. 2B, Fig. S2B**).

**Fig. 2.**
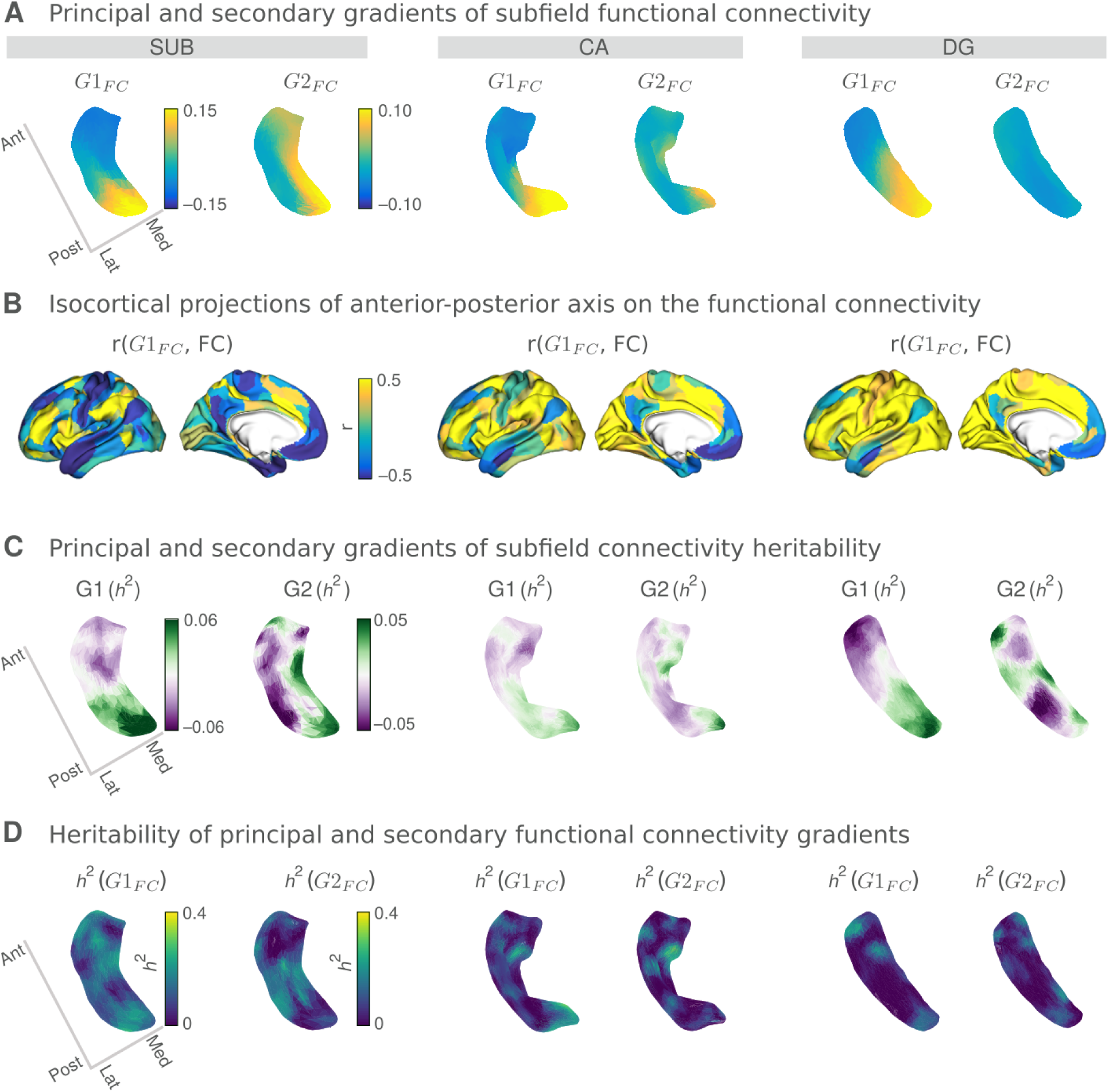
Topological representations of hippocampal functional organisation and their heritability. **A**. Connectivity gradients of subfield-isocortical FC for SUB (left), CA (middle) and DG (right). Gradient (*G*1_*FC*_) depicts an anterior-posterior (A-P) connectivity axis, whereas Gradient 2 (*G*2_*FC*_) displays a medial-lateral axis. **B**. Variations in hippocampal-isocortical FC across the projected on the isocortex (Pearson’s r-values). Lower r-values (blue) indicate FC similarity between the anterior subfield portions and isocortex, whereas higher r-values (yellow) that of the posterior subfield portions and isocortex. **C**. Gradient profiles of hippocampal FC heritability. Gradient 1 of the FC heritability (G1 (*h*^2^)) depicts an A-P separation of the *h*^2^ profiles for all subfields. **D**. Heritability scores of subfield FC gradients (*h*^2^ (*G*1_*FC*_) and *h*^2^ (*G*2_*FC*_)).

Next, we decomposed the heritability scores of the hippocampal-isocortical connectome to probe a potential organisational axis underlying the heritability of individual FC measures (**Fig. 2C**). The primary gradient of FC heritability G1 (*h*^2^) depicted an A-P separation of the *h*^2^ profiles for all the subfields. The secondary gradient of FC heritability G2 (*h*^2^) traversed the M-L axis for SUB but did not reveal a clear pattern for CA and DG. We further obtained the heritability of the A-P and M-L functional gradients themselves, as represented in **Fig. 2A**, *h* ^2^ (*G*1_*FC*_) and *h*^2^ (*G*2_*FC*_), respectively, to assess whether individual variations in *local* gradient loadings were heritable (**Fig. 2D**). For all subfields *h* ^2^ (*G*1_*FC*_) was found to be modest to low (SUB: mean: 0.14, range: [0, 0.29]; CA: mean: 0.08, range: [0, 0.32]; DG mean: 0.05, range: [0, 0.27]). Also the second gradient’s heritability, *h* ^2^ (*G*2_*FC*_), was found to be modest to low for all subfields (SUB: mean: 0.09, range: [0, 0.30], CA: mean: 0.06, range: [0, 0.39], and DG: mean: 0.06, range: [0, 0.23]). The heritability strength of both functional gradients did not show a clear spatial pattern (**Fig. S2C)**.

### Hippocampal microstructure is highly heritable and shows genetic correlation with the isocortex along its intrinsic organisational axes (Figure 3)

Having shown that functional connectivity of hippocampal subfields is heritable, but intrinsic functional organisation of subfields is less so, we aimed to evaluate the heritability of hippocampal subfield structure. To do so, we utilised individual T1w/T2w intensity maps to probe microstructure *in vivo* (**Fig. S3**). Local T1w/T2w maps were highly heritable across all subfields, reaching up to *h*^2^ = 0.77 for SUB (mean ± SD = 0.44 ± 0.15 for SUB, 0.41 ± 0.12 for CA, and 0.43 ± 0.07 for DG) (**Fig. 3A**). Multiple comparison corrections using FDR reported significant heritability scores across almost all subfield vertices. By adjusting for the mean T1w/T2w as a covariate in the heritability model, we found similar heritability patterns for individual subfields, and both hemispheres (**Fig. S4**). This indicates that the heritability of subfields was present beyond any mean T1w/T2w intensity variation across individuals.

**Fig. 3.**
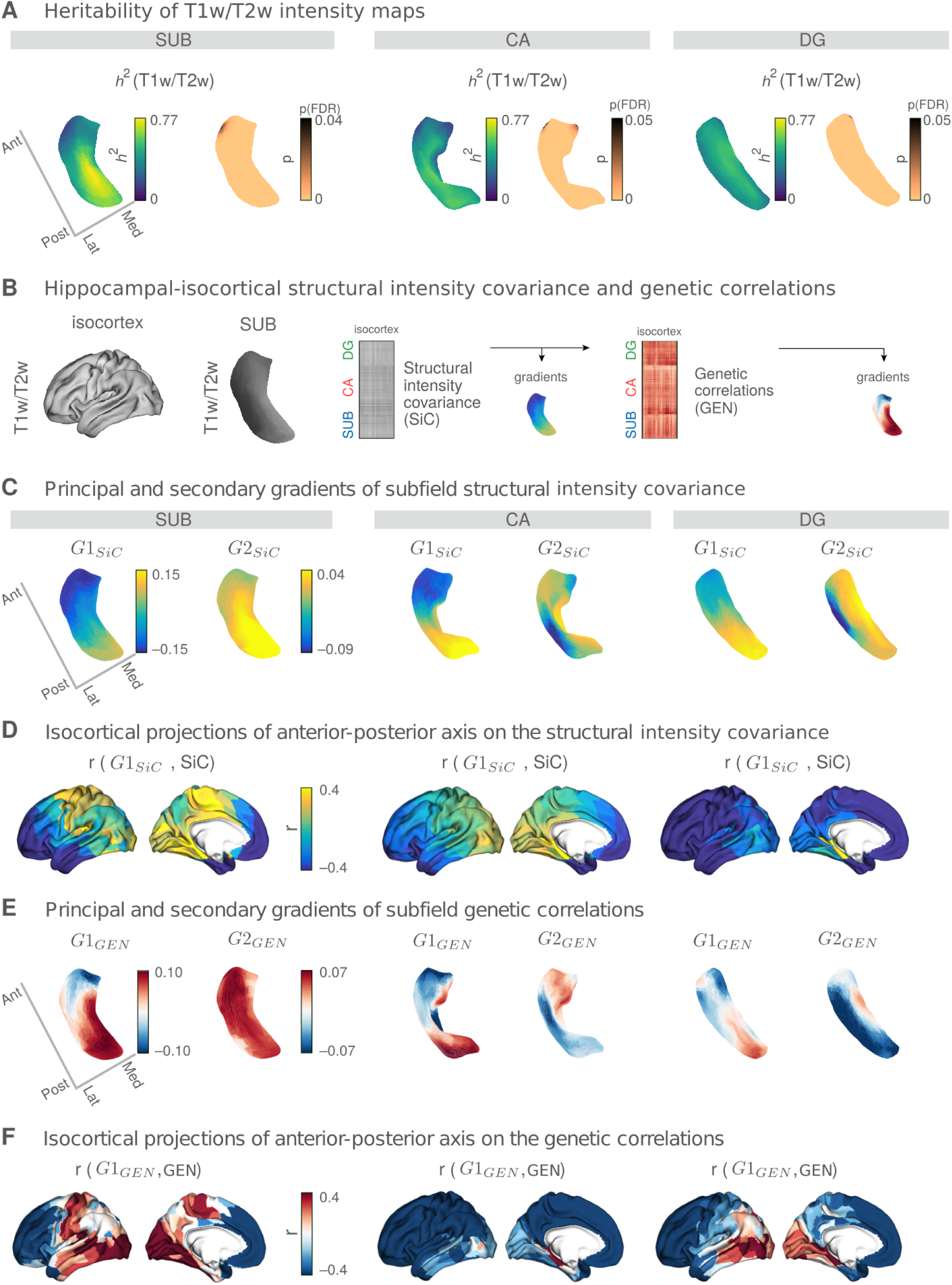
Hippocampal microstructural organisation and its heritability. **A**. Heritability of subfield T1w/T2w profiles (*h* ^2^(T1w/T2w)) and its significance levels. T1w/T2w maps were strongly heritable across all subfields. p-values were reported after multiple comparison corrections using FDR (copper colour denotes pFDR < 0.05, black pFDR > 0.05). **B**. Hippocampal-isocortical structural intensity covariance (SiC) was assessed by correlating hippocampal and isocortical T1w/T2w intensity maps across participants and subfields. Shared genetic variations in T1w/T2w intensity maps were assessed by conducting a genetic correlation (GEN) analysis on the SiC. Both SiC and GEN matrices were then decomposed into their gradient representations, separately. **C**. Gradients of SiC for SUB (left), CA (middle), and DG (right). *G*1_*SiC*_ represents an anterior-posterior (A-P) axis for all subfields, whereas *G*2_*SiC*_ reflects the differential axis of local transitions for individual subfields. **D**. Variations in SiC across its *G*1_*SiC*_ projected on the isocortex (Pearson’s r-values). Lower r-values (blue) indicate SiC similarity between the anterior subfield portions and isocortex, whereas higher r-values (yellow) that of the posterior subfield portions and isocortex. **E**. Gradients of GEN for SUB (left), CA (middle), and DG (right). *G*1_*GEN*_ represents an A-P axis for all subfields, whereas *G*2_*GEN*_ reflects the differential axis of local transitions for individual subfields. **F**. Variations in GEN across its *G*1_*GEN*_ projected on the isocortex (Pearson’s r-values). Lower r-values (dark blue) depict shared genetic influence between anterior subfield portions and isocortex and higher r-values (red) that of posterior subfield portions and isocortex.

To evaluate the spatial similarity between local microstructure and functional gradients, we quantified the group-level association between the T1w/T2w and *G*2_*FC*_ for subfields (**Fig. S5A**). T1w/T2w maps had the highest correlation with *G*2_*FC*_ for the SUB (Pearson’s r = 0.93, p-value after spatial autocorrelation correction ^56^; *p* _*vario*_ < 0.001), and less with the other subfields (CA: r = 0.23, *p* _*vario*_ = 0.02, DG:r = -0.01, *p* _*vario*_ = 0.9). Furthermore, individual-level T1w/T2w and *G*2_*FC*_ correlations were found to be significantly positive across participants for SUB (median 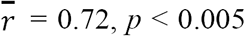, one-tailed Wilcoxon signed-rank test) and CA 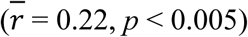, however not for DG (**Fig. S5B**).

Then, we evaluated whether there is a genetic correlation between microstructure of hippocampal subfields and that of the isocortex, to probe whether potential co-variation of hippocampal and isocortical microstructure is governed by shared genetic factors. We first correlated vertex-wise subfield and parcel-wise isocortical T1w/T2w maps across all participants (n = 709), resulting in a structural intensity covariance (SiC) matrix (**Fig. 3B**). Using gradient decomposition, we evaluated intrinsic axes of covariance/genetic correlation within subfields based on their correspondence with isocortical microstructure ^57^. The principal gradient of SiC (*G*1_*SiC*_) revealed an A-P organisational axis across all the subfields **(Fig. 3C)**. We observed a high similarity between *G*1_*SiC*_ and *G*1_*FC*_ profiles (SUB: r = 0.88, CA: r =0.86, and DG: r =0.88, *p* < 0.001 for all subfields). The second gradient of SiC (*G*2_*SiC*_) did not represent a converging organisational pattern for the subfields. Evaluating the pattern of correlation between subfield-isocortex SiC and *G*1_*SiC*_, we could assess how hippocampal and isocortical regions spatially relate to each other in terms of their microstructural similarity (**Fig. 3D**). Anterior hippocampal portions (blue in **Fig. 3C**) shared more microstructural similarity with the anterior isocortex, in particular anterior frontal and temporal cortex, while the posterior hippocampal portions (yellow in **Fig. 3C**) were related to visual cortex, lingual and fusiform areas, and sensorimotor cortex for SUB, and visual cortex, lingual and fusiform areas for CA. For the DG, we observed less divergent patterns of subfield-isocortical similarity between its anterior and posterior portions, with anterior portions relating to all of the isocortex except for visual, lingual and fusiform areas which showed a positive relation to posterior parts of DG.

Following, we computed genetic correlation (GEN) between subfields and isocortex ^55^. A high GEN score indicates that the microstructural phenotype of subfield and isocortex are influenced by the same set of genes ^58^. The principal gradient of the GEN (*G*1) again displayed an A-P axis for all the subfields (**Fig. 3E**). Also *G*1_*GEN*_ showed spatial similarity with *G*_*FC*_ (SUB: r = 0.67, CA: r = 0.41, DG: r = 0.75, and *p*_*vario*_ < 0.001 for all subfields). The second gradient (*G*2_*GEN*_) did not reveal a consistent organisational axis within each subfield, but rather varied between subfields. Analogous to SiC, we then investigated the correlation between hippocampal-isocortical GEN variations and (**Fig. 3F**). Indeed, patterns were largely mirroring those observed in SiC, indicating that the structural intensity covariance between hippocampus and isocortex is largely concordant with genetic patterning.

### Coupling of hippocampal subfield function and microstructure (Figure 4)

Last, we studied the shared organisation of hippocampal function and structure to evaluate whether regions with similar microstructure in hippocampus and isocortex also show a functional connection as predicted by the structural model. To do so, we computed the coupling of microstructure covariance and functional connectivity between the subfield and isocortex at each vertex of the subfields. Second, to probe whether the similarity of microstructure and functional profiles varied along the respective subfields’ intrinsic functional and structural axes, we computed the degree of hippocampal organisational axes’ similarity using the coefficient of determination (*R*^2^) (**Fig. 4**). For SUB, we found a dominant pattern of A-P axes shared by *G*1_*FC*_, *G*1_*SiC*_, *G*1_*GEN*_ and *G*2_*SiC*_. However, the M-L axes, reflecting variation in local T1w/T2w and *G*2_*FC*_, best described the coupling between microstructure and function (*R*^2^ = 0.35), with lateral regions showing moderately positive coupling and medial regions showing low coupling. For CA, coupling rather followed a posterior (high) to anterior (low) pattern, corresponding to *G*1_*FC*_, *G*1_*SiC*_, *G*1_*GEN*_ and *G*2_*GEN*_ (0.33 < *R*^2^ < 0.64). Last, for DG we found moderate variation in coupling, which showed a spatial relation to *G*1_*FC*_, *G*1_*SiC*_, *G*1_*GEN*_ and *G*2_*GEN*_ (0.30 < *R*^2^ < 0.49). Here, posterior regions showing increased and anterior regions showing decreased coupling between microstructure covariance and functional connectivity.

**Fig. 4.**
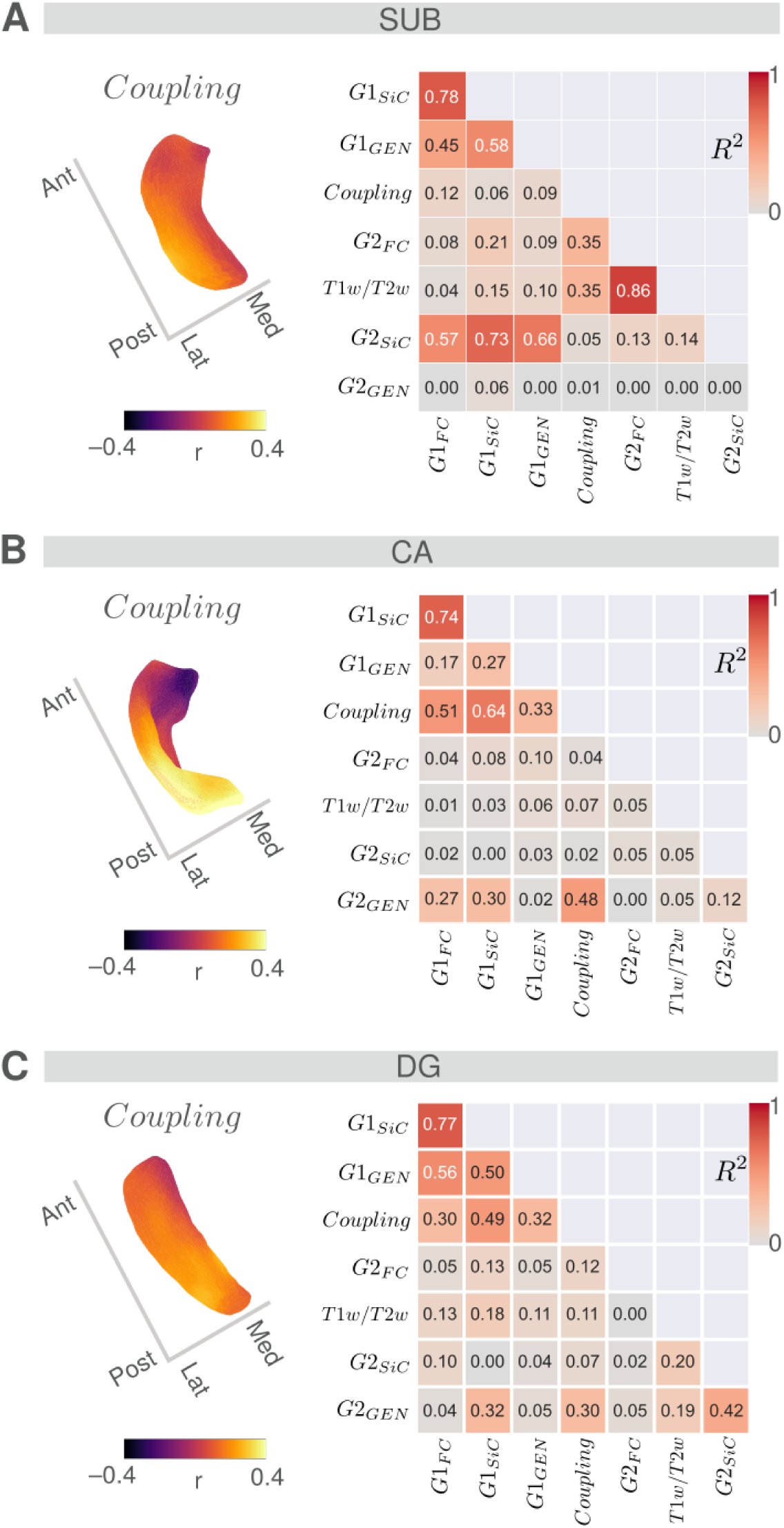
Hippocampal structural-functional coupling maps and associations among organisational axes. **A. SUB:** Subfield vertex-wise coupling map between hippocampal-isocortical functional connectivity (FC) and structural intensity covariance (SiC) (upper panel). Higher coupling values (Pearson’s r) denote an association between FC and SiC, whereas lower coupling values display a dissociation between them. Spatial similarity between hippocampal organisational axes (*G*1 − *G*2_*FC*_, *G*1 − *G*2_*SiC*_, *G*1 − *G*2_*GEN*_, coupling and T1w/T2w maps) is denoted by the coefficient of determination *R*^2^. High *R*^2^ values (red) indicate a strong spatial alignment between the organisational axes, whereas low *R*^2^ values (grey) an unalignment. Panels **B**. and **C**. display coupling maps and *R*^2^ values for **CA** and **DG**, respectively.

## Discussion

The hippocampus is a densely interconnected region where stability and plasticity coincide. Building on emerging work describing the coalescence of anterior-to-posterior and medial-to-lateral gradients of function and microstructure in the hippocampal formation *in vivo* ^21,35,59,60^, we describe the heritability of hippocampal functional and structural organisation as well as its genetic relationship with the isocortex. First, we found that functional connectivity of hippocampal subfields to isocortex was heritable. However, intrinsic functional organisation of hippocampal subfields showed only marginal heritability. At the same time, we found that spatial variations in subfield T1w/T2w intensity maps, serving as a marker for myelin-related microstructure ^50^, were heritable and topographically related to local subfield functional organisation. Exploring the covariance of local hippocampal and isocortex microstructure, we found they consistently followed an A-P axis, with anterior subfield regions relating to anterior frontal and temporal cortex, whereas posterior subfield regions relating to visual and inferior temporal areas. These patterns were genetically correlated, indicating that microstructure of the subfields underlie shared genetic influences with the isocortex. Last, evaluating the similarity by the structure-function coupling in the hippocampal subfields along intrinsic hippocampal axes, we found lateral/posterior regions to be highly coupled, whereas anterior/medial regions were uncoupled. This illustrates how genetic axes may scaffold hippocampal structure and function, enabling a decoupling of function from structure in particular in anterior/medial areas, genetically linked to anterior/transmodal portions of the isocortex.

To study the heritability of subregional functional and microstructural organisation of the hippocampal formation, we automatically segmented the hippocampal formation via a subfield and surface-based approach (SUB, CA, and DG) ^52^, which has been previously validated in both healthy individuals and those with hippocampal pathology ^11^. Such surface-based approaches improve anatomical alignment across individuals ^59^. In the current work, we could replicate previously established hippocampal-cortical FC organisation across subfields. The primary gradient demonstrated A-P transitions (long axis ^22,61–63^), whereas the secondary gradient revealed M-L separations (transverse axis ^21,27,64^). Second, as in previous work, the M-L axis was found to align strongly with the microstructural proxy, particularly for the SUB and to a lesser extent in CA. In sum, we could replicate previous work ^35^ and again observe that specialisation of the long axis was preserved in all subfields, whereas the transverse axis indicated a link between intrinsic FC and microstructure, particularly in SUB.

Extending previous work describing mean axes of microstructural and functional organisation of hippocampal subfields, we investigated whether individual variation in hippocampal organisation was partly attributable to genetic factors. Previously, volume-based studies segmented the hippocampus into 12 subregions and showed that these volumes exhibit strong heritability ^38,65^. Moreover, genome-wide studies identified single-nucleotide polymorphisms (SNP) associated with hippocampal volumes ^66–68^ showing, in part, unique SNPs for each subfield ^69^. Here, we extended this work by studying the heritability of subtle variations of microstructure and function within subfield surfaces, as well as their link to the isocortex. We observed highest heritability within the subfield microstructure proxy (T1w/T2w) and lowest for the A-P and M-L functional hippocampal gradients. Indeed, the heritability of both functional gradients was moderate to low, indicating that individual variation within functional gradients did not vary strongly as a function of genetic proximity of individuals. At the same time, we found that heritability of subfield-isocortical FC was again organised along an A-P axis, indicating that anterior and posterior portions of hippocampal subfields have distinct and heritable relations with the isocortex. Moreover, we found that the functional M-L gradient correlated strongly with T1w/T2w microstructure along the hippocampal subfield surfaces, which, in turn, was highly heritable. It is possible that the intrinsic, heritable, structural axes within the hippocampus scaffold a more flexible functional organisation. Indeed, environmentally induced brain changes may be interpreted as a degree of aberration from the heritability, i.e. the less heritable a brain region/metric, the larger the potential environmental influence ^43,70,71^. It is thus possible that the low heritability of functional organisation of subfields reflects these variations, attributable to environmental effects and associated with hippocampal plasticity. Indeed, previous work in rodents and human adults has shown high plasticity of the hippocampus in both species ^72^, which has been linked to internal and external changes, such as hormonal levels and stress responses ^40^.

As the internal wiring of the hippocampus relates to its connectivity to the rest of the brain, we evaluated the genetic relationship between subfield surface microstructure and isocortical microstructure. To do so, we probed the covariance between hippocampal subfields and isocortex microstructure (structural intensity covariance, SiC), and their genetic correlation. SiC emphasises the morphological similarity among brain regions, with high covariance between two regions across individuals indicating these regions share maturational and genetic trajectories ^73^. Although the decomposed SiC and genetic correlation measure originated from the T1w/T2w maps, its low dimensional components depicted similar spatial organisation to that of the functional maps. The primary covariance gradient revealed an A-P axis for all the subfields, which was mirrored by a highly similar gradient based only on the genetic correlation between local subfield and isocortical microstructure. This indicates a distinction between microstructure of anterior and posterior regions of hippocampal subfields based on its genetic similarity with the isocortex, which was found to be mirrored in its functional organisation in the current sample. Regions in anterior parts of the subfields showed a genetic similarity with anterior frontal and temporal cortex, whereas those in posterior parts of the subfields showed a genetic similarity with posterior occipital-temporal regions. Earlier studies have presented an isocortex-wide A-P topography derived from cortical thickness morphology ^74^, microstructural profile covariance ^46^, and grey matter volumes ^25^. The isocortical A-P topography resembles a frontal-polar differentiation of myelin density ^75,76^ and shows spatial similarity with a cortical functional gradient traversing between the transmodal to unimodal axis ^76,77^. In line with our observation that morphometric similarity of hippocampus and isocortex is genetically determined ^78,79^, the concordance of genetic similarity between the A-P subfield axis and A-P isocortical axis has been previously reported using transcriptomic data ^36^. Thus, the internal, heritable, organisation of hippocampal subfield microstructure has a genetic correlation with isocortex, which spatially co-varies with its functional organisation.

Beyond similarities, we also observed differences in subfield-isocortical genetic associations, both in the primary and secondary covariance and genetic correlation gradient. For example, we found a clear differentiation between genetic relationships of hippocampus and isocortex along the anterior-posterior axes with posterior regions of CA, showing only little association with temporal-occipital regions, and SUB showing a clear distinction between anterior subfield regions and its correspondence to anterior frontal/temporal cortex and posterior subfield regions and its correspondence to temporo-occipital and sensory regions. Moreover, the second genetic correlation gradient varied strongly between SUB, CA, and DG, suggesting to vary rather as a function of subfield infolding. This may relate to the subfield specific neurodevelopmental trajectories. For example, the CA - Ammon’s Horn - is one of the first brain regions to develop in the prenatal period ^38,80^. Conversely, the SUB extends its maturation towards the postnatal period ^81^. Finally, DG maturation exceeds the postnatal period ^82^, possibly underscoring posterior parietal associations. Thus, timing of pre- and post-natal development may be reflected in the genetic similarity patterning between subfields and their association with the isocortex.

Lastly, to understand whether hippocampal function and structure covary along genetic organisational axes, we assessed local coupling maps of structural and functional subfield-isocortical profiles. Evaluating differences between structural and functional organisation, we found that coupling between structure and function was highest in posterior/medial portions of the hippocampal subfields, whereas anterior portions were uncoupled. Moreover, the coupling of hippocampal microstructure and function shows covariation with intrinsic functional and structural axes that we showed to align with genetic correlation to the isocortex. In particular, in the hippocampus, we found that posterior regions have a predominant structural and functional association with unimodal cortical regions whereas anterior regions are linked to transmodal cortex, similar to previous reports ^36,60^. Thus, mirroring observations in the cortex ^43,45^, it may be that portions of the hippocampus associated with posterior/unimodal regions show more similarity between structure and function than those related to transmodal areas such as anterior frontal and temporal cortex. Functionally, the anterior hippocampus has been reported to participate in associative memory processing ^83^, in which DMN is also involved and known to be integrating with parietal and temporal lobes for episodic memory retrieval ^84^. Conversely, the posterior hippocampus is suggested to be a mediator for spatial memory encoding ^85^, in which parietal cortices ^86^ and attention and salience networks are recruited ^87^. Together, the divergence observed in function and structure along genetic axes of hippocampal subfields may reflect a hierarchy of complexity, with more uncoupled portions of the hippocampus enabling more flexible forms for cognitive processing, an important hypothesis for future studies to examine.

In sum, we showed that hippocampal subfields are organised along heritable posterior-to-anterior and medial-to-lateral axes which show a genetic link to isocortical functional and structural organisation. Future work may evaluate the association between maturational axes in cortical structure and divergent functional profiles along the hippocampal formation. This may provide an important step to better understand how the anatomy of the hippocampus supports its unique and versatile function.

## Materials and Methods

### Participants

Participants from the HCP S900 release ^49^ with four complete resting-state fMRI sessions and high-resolution structural images were selected. Participant recruitment procedures and informed consent forms, including consent to share de-identified data, were previously approved by the Washington University Institutional Review Board as part of the HCP. Participants with anatomical anomalies or tissue segmentation errors listed in the HCP issues were discarded (n = 40). For every participant, we segmented hippocampal subfields: subiculum (SUB), CA1-3 (CA), and CA4-DG (DG) along the structural images using a patch-based surface algorithm ^52^. Based on a visual inspection of subfield delineations, we discarded participants with poor segmentation quality (n = 42). As a necessity for the functional connectivity (FC) gradient analysis (see section **Functional Connectivity and Gradients**), we further excluded participants (n = 31), whose FC maps were poorly associated with the group-level reference FC. There remained n = 709 participants (395 women, mean ± SD age = 28.7 ± 3.7 y) accessible for our study. Among the 709 participants included in this study, there were 176 monozygotic twins, 178 siblings without twin status and 355 participants without familial relatedness. All quality assessment steps and analysis scripts used in this study are available at https://github.com/CNG-LAB/cngopen/tree/main/hippocampus.

### Neuroimaging Data Acquisition and Preprocessing

Details of the HCP neuroimaging protocol and processing pipelines are available at ^88^. In brief, we extracted T1-weighted (T1w) and T2-weighted (T2w) images available in the HCP initiative, which were all acquired on a 3T Siemens Skyra scanner. T1w images were acquired using a three-dimensional magnetization prepared rapid gradient-echo (3D-MPRAGE) sequence (0.7 mm isotropic voxels, matrix = 320 × 320, 256 sagittal slices, TR = 2400 ms, TE = 2.14 ms, TI = 1000 ms, flip angle = 8°, iPAT = 2). Resting-state fMRI images were acquired using a multi-band accelerated 2D-BOLD echo-planar imaging (EPI) sequence (2 mm isotropic voxels, matrix = 104 × 90, 72 sagittal slices, TR = 720 ms, TE = 33 ms, flip angle = 52°, mb factor = 8, 1200 volumes/scan). The fMRI data was collected at two sessions (1, 2) and in two phase encoding directions at each session (left-right [LR] and right-left [RL]), resulting in four resting-state fMRI datasets in total ([LR1], [RL1], [LR2], [RL2]).

Preprocessing steps for the structural MRI images included gradient nonlinearity correction, brain extraction, distortion correction and co-registration of T1w and T2w images using rigid body transformations. Then, an intensity nonuniformity correction was performed using T1w and T2w contrasts ^50^ and subcortical structures were segmented using FSL FIRST ^89^. Subsequently, preprocessed images were nonlinearly registered to the MNI152 template and cortical surfaces were reconstructed with FreeSurfer 5.3.0-HCP ^90–92^. Finally, the individual cortical surfaces were registered to the Conte69 template ^93^ using MSMA11 ^53^.

Preprocessing of rs-fMRI images included corrections for the gradient nonlinearity, head motion and distortion. The images were then aligned to the T1w space using rigid-body and boundary-based registrations together ^94^. The transformation matrices from this alignment step and that of the earlier T2w to T1w alignment were concatenated and applied to the rs-fMRI images at a single interpolation step to warp rs-fMRI images to the MNI152. Further processing in MNI152 space included bias field removal, whole brain intensity normalisation, high pass filtering (> 2000s FWHM) and noise removal with the ICA-FIX procedure ^95^.

### Hippocampus Subfield Segmentations

We used the SurfPatch algorithm ^52^ to automatically delineate the hippocampal subfields of all participants: subiculum (SUB), CA1-3 (CA), and CA4-DG (DG). SurfPatch is a multi-template surface-path hippocampal segmentation method trained on a public dataset of manual segmentations in healthy controls ^96^, and has been validated in patients with histopathology of the hippocampus ^11^. This algorithm incorporates a spherical harmonic shape parameterization and point distribution model of the surfaces ^97^. Next, to minimise partial volume effects, we generated medial surfaces running through the centre of each subfield using a Hamilton-Jakobian approach ^98^. The spherical harmonic parameterization was propagated to the medial sheet to improve vertex-correspondence across individuals based on shape inherent information. Resultant CA surfaces consisted of 10242 vertices and both DG and SUB surfaces of 5762 vertices. Next, CA, DG, and SUB surfaces were further downsampled to 2048, 1024, and 1024 vertices, respectively. All surface segmentations underwent a visual inspection and are available upon request.

### Isocortex and Subfield Time Series

We mapped medial sheet meshes and volumetric resting-state fMRI data to native T1w space. Time series were sampled at each hippocampal and cortical mid-thickness vertex ^53^. Hippocampal surface features were smoothed using a Gaussian diffusion kernel with 5 mesh units as FWHM in all subfields and isocortex. Sampling was carried out in a native T1w space to minimise the interpolation. Cortical time series were averaged within a previously established multi-modal parcellation scheme of the Glasser Atlas of 360 areas (180 regions per hemisphere) ^53^. Surface-based time series were smoothed using a Gaussian diffusion kernel with 5 mesh units as full-width-at-half-maximum (FWHM).

### Functional Connectivity

For every participant separately (n = 740), we computed the linear correlation coefficients between isocortex-wide time series (360×1200) and hippocampal subfield time series for SUB (1024×1200), CA (2048×1200), and DG (1024×1200). This resulted in a isocortex wide functional connectivity (FC) map (360×1) for every subject and subfield. We obtained group-level reference FC maps for every subfield by averaging individual FC maps across participants. We further profiled the similarity of individual FC maps to the reference FC maps by means of simple correlation (**Supplementary Fig. S1A**). Participants with a lower degree of similarity (r < 0.45) to the reference map were excluded (n = 31). Finally, the FC map of the isocortex to each hippocampal subfield for the remaining 709 participants was mapped using linear and mixed effects models in BrainStat (https://github.com/MICA-MNI/BrainStat).

### T1w/T2w Maps and Structural Intensity Covariance

To study microstructural features of the hippocampus, we used the ratio of T1-over T2-weighted (T1w/T2w) image intensities. We resampled native T1w/T2w images to the MNI152 space and mapped them to hippocampal subfield surfaces (SUB, CA, DG) using Connectome Workbench (v1.4.2, volume-warpfield-resample and volume-to-surface-mapping tools) ^99^. To assess the quality of T1w/T2w intensities projected on the hippocampal subfields, we obtained the mean T1w/T2w intensity distributions of all participants for potential outlier detection **(Fig. 2A, Supplementary Fig. S3C)**. We computed the structural intensity covariance (SiC) by correlating hippocampal and cortical T1w/T2w intensity maps resulting in 1384×1384 matrix for SUB, 2408×2408 matrix for CA, and 1384×1384 matrix for DG.

### Heritability and Genetic Correlation

Heritability and genetic correlation analysis were conducted with the Sequential Oligogenic Linkage Analysis Routines (SOLAR, v8.5.1, http://www.solar-eclipse-genetics.org/). SOLAR employs a maximum likelihood variance-decomposition approach optimised to perform genetic analyses in pedigrees of arbitrary size and complexity ^55,100^. SOLAR models genetic proximity by covariance between family members ^55,100^.

In brief, heritability (i.e. narrow-sense heritability *h*^2^) is defined as the proportion of the phenotypic variance 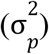 in a trait that is attributable to the additive effects of genes 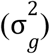, i.e. 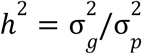. SOLAR estimates heritability by comparing the observed phenotypic covariance matrix with the covariance matrix predicted by kinship ^55,100^. Significance of the heritability estimate was tested using a likelihood ratio test where the likelihood of a restricted model (with 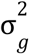 constrained to zero) is compared with the likelihood of the estimated model. Twice the difference between the log likelihoods of these models yields a test statistic, which is asymptotically distributed as a 50:50 mixture of a χ^2^ variable with 1 degree-of-freedom and a point mass at zero ^55,100^. We quantified the heritability of (i) hippocampal-isocortical functional connectivity patterns, (ii) hippocampal subfield gradients, and (iii) T1w/T2w intensity maps. We included covariates in all heritability analyses including. *age, sex, age* × *sex, age*^2^ and *age*^2^ × *sex*.

To estimate if variations in T1w/T2w intensity maps between hippocampus and isocortex were influenced by the same genetic factors, a genetic correlation analysis was conducted. Genetic correlations indicate the proportion of variance that determines the extent to which genetic influences on one trait are shared with genetic influences on another trait (e.g. pleiotropy). In SOLAR, the phenotypic correlation (ρ_*p*_) was decomposed through bivariate polygenic analyses to estimate genetic (ρ_*g*_) and environmental (ρ_*e*_) correlations using the following formula: 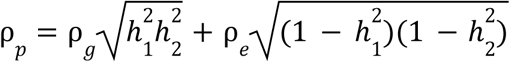, where 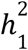 and 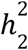 are the heritability estimates of the vertex-based values in hippocampus and isocortex ^58,101^. The significance of these correlations was determined (similar to heritability analyses) by likelihood ratio tests comparing a model in which ρ_*g*_ was estimated with a model in which ρ_*g*_ was constrained to zero (no shared genetic effect) and constrained to 1 (complete pleiotropy) ^58,101^.

### Connectivity Gradients

Using the diffusion embedding algorithm, we generated low-dimensional representations of hippocampal-cortical FC, namely the *gradients*. For every participant separately (n = 709), we computed the linear correlation coefficients between isocortex-wide time series 360 × 1200 and hippocampal subfield time series for SUB (1024 × 1200), CA (2048 × 1200), and DG (1024 × 1200) to identify the FC matrices. We computed hippocampal subfield FC gradients, similarly to those identified by Vos de Wael (2018). Next, the time series of each vertex was correlated with the isocortex wide time series, yielding a hippocampal-cortical FC map (4096×360) for every subject. We used BrainSpace ^35,57^ to derive connectivity gradients from the group-level FC matrix using diffusion map embedding (normalised angle kernel, 90th percentile thresholding for the sparsity, and diffusion time estimation of α = 0.5) ^51^.

Along each single gradient (4096×1), hippocampus vertices that share similar connectivity patterns have similar embedding values. The first and second gradients explained 32.4% of the total variance in the subfield FC map (**Supplementary Fig. S1B, S1C**). Having validated the gradient representations of hippocampal subfields at the group-level, we computed the individual-level gradients for every participant. Subsequently, individual gradients were aligned to the group-level gradients using Procrustes alignment to be scaled onto a common embedded connectivity space.

## Acknowledgements

We would like to thank the various contributors to the open access databases that our data was downloaded from. Funding: HCP data were provided by the Human Connectome Project, Washington University, the University of Minnesota, and Oxford University Consortium (Principal Investigators: David Van Essen and Kamil Ugurbil; 1U54MH091657) funded by the 16 NIH Institutes and Centers that support the NIH Blueprint for Neuroscience Research; and by the McDonnell Center for Systems Neuroscience at Washington University. This study was supported by the Deutsche Forschungsgemeinschaft (DFG, EI 816/21-1), the National Institute of Mental Health (R01-MH074457), the Helmholtz Portfolio Theme “Supercomputing and Modeling for the Human Brain’’ and the European Union’s Horizon 2020 Research and Innovation Program under Grant Agreement No. 785907 (HBP SGA2). R.V. was supported by the Richard and Ann Sievers award. S.L.V. was supported by Max Planck Gesellschaft (Otto Hahn award). B.C.B. acknowledges support from the SickKids Foundation (NI17-039), the National Sciences and Engineering Research Council of Canada (NSERC; Discovery-1304413), CIHR (FDN154298), Azrieli Center for Autism Research (ACAR), an MNI-Cambridge collaboration grant, and the Canada Research Chairs program. Last, this work was funded in part by Helmholtz Association’s Initiative and Networking Fund under the Helmholtz International Lab grant agreement InterLabs-0015, and the Canada First Research Excellence Fund (CFREF Competition 2, 2015-2016) awarded to the Healthy Brains, Healthy Lives initiative at McGill University, through the Helmholtz International BigBrain Analytics and Learning Laboratory (HIBALL). M.D.H. was funded by the Max Planck Society and the German Ministry of Education and Research.

## Author contributions

Ş.B., and S.L.V. conceived and designed the analysis, performed the analysis, wrote the draft manuscript and revised the manuscript. R.V. and H.L.S. aided in data analysis. A.B., B.C., N.B, R.V., and B.C.B. provided hippocampal subfield segmentations. All authors helped with writing and revising the manuscript.

## Competing interests

The authors declare no conflict of interest.

## Supplementary Materials

**Supplementary Fig. S1.**
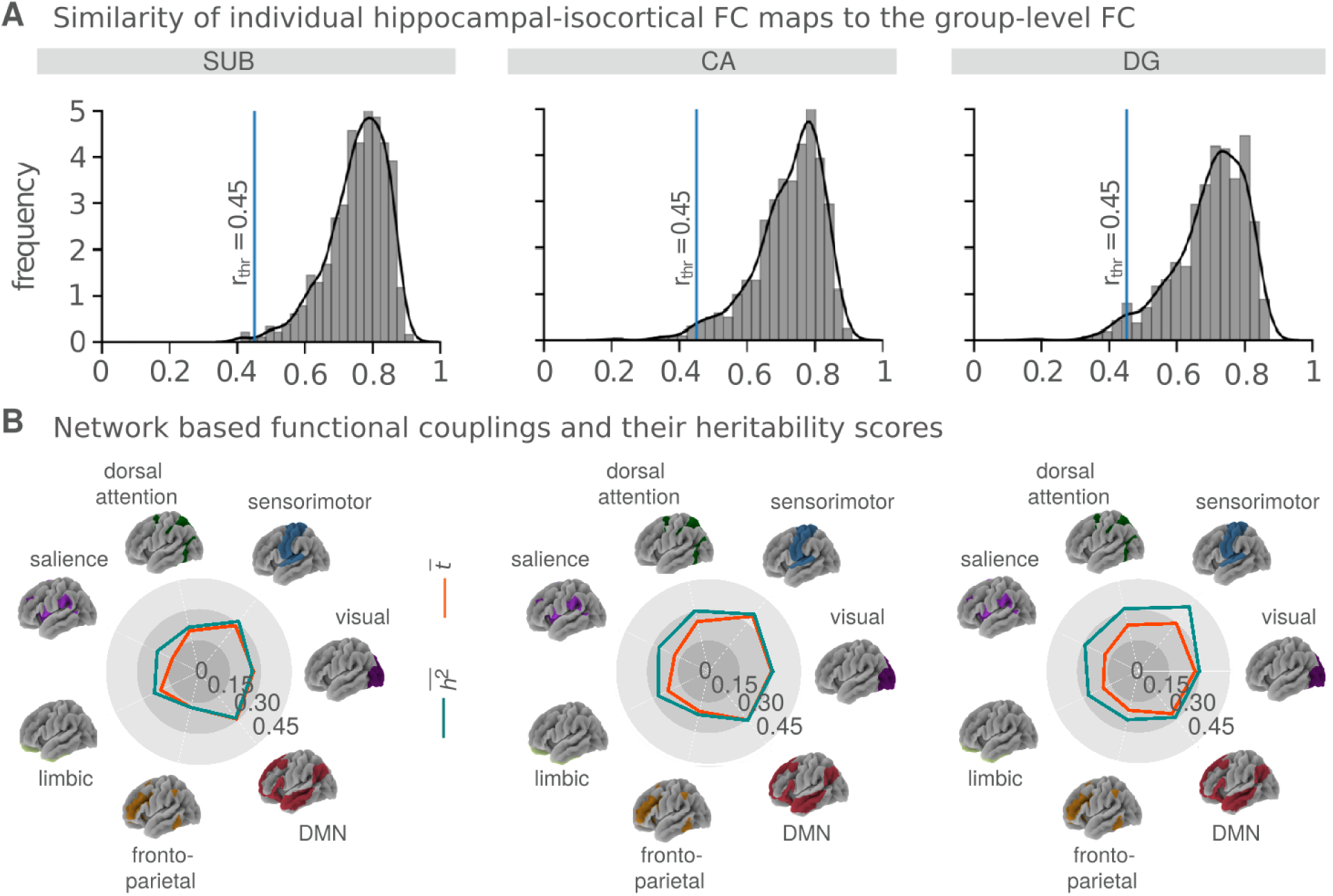
Quality assessment for the individual hippocampal-isocortical functional connectivity (FC) profiles and network-level representations of the group-level hippocampal-isocortical FC. **A**. FC similarity of n = 740 participants to the group-level FC quantified by means of Pearson’s correlations (r). Threshold r-value (*r*_*thr*_ = 0.45) was assessed by computing the 2.5 standard deviation distance away from the mean r-values. Participants with a lower degree of similarity (*r*_*thr*_ < 0.45, n = 31) were excluded prior to the functional connectome gradient analysis. **B**. Hippocampal-isocortical FC strength (t-values) and its heritability (*h*^2^scores) distributed into seven networks ^102^. The t-values and *h*^2^ scores were averaged (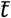 and 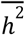, red and blue, respectively) within each single network and demonstrated with a spyder plot.

**Supplementary Fig. S2.**
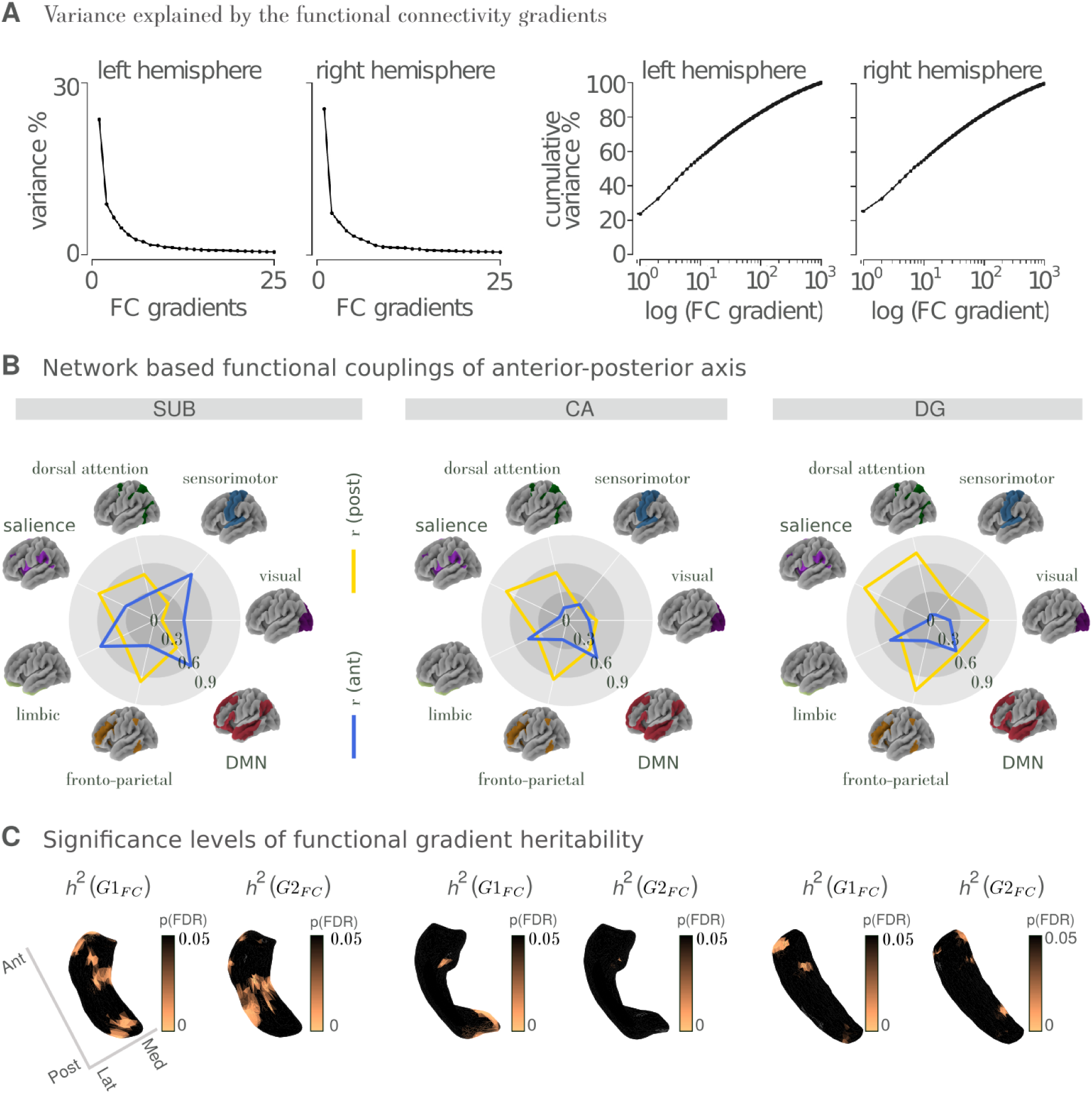
Connectivity gradients of hippocampal-isocortical functional connectivity (FC) and their heritability. **A**. Variance explained by the functional connectivity gradients. The first gradients explained 23.6% of total hippocampal-isocortical FC variance for the left hemisphere and 25.5% for the right hemisphere. Cumulative variance explained by the first three gradients was 38.5% for the left hemisphere and 38.9% for the right hemisphere. **B**. FC from anterior (r-values, blue) and posterior (r-values, yellow) subfield portions revealed by the *G*1_*FC*_ were distributed into seven large-scale functional networks ^102^. **C**. Significance levels of gradient map heritability (*h*^2^(*G*1_*FC*_) and *h* ^2^(*G*2_*FC*_)) reported by p(FDR).

**Supplementary Fig. S3.**
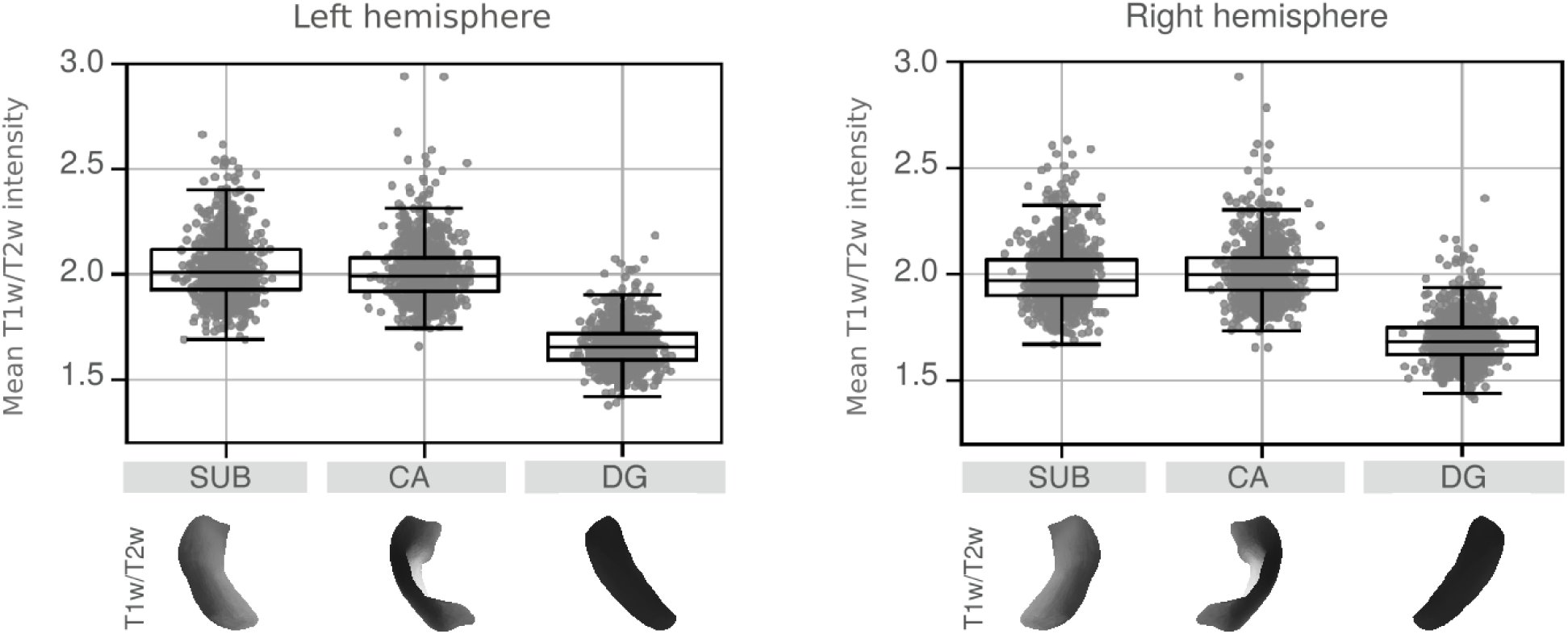
Quality assessment on individual (n = 709) T1w/T2w intensity maps for left and right hemispheres. Mean T1w/T2w intensity map distributions for each subject (n = 709, x-axis) and each subfield (SUB, CA, DG, y-axis) are depicted for left and right hemispheres, separately. For the left hemisphere, mean and standard deviations of T1w/T2w maps were 2.02 ± 0.41 for SUB, 2.01 ± 0.78 for CA, and 1.66 ± 0.22 for DG. For the right hemisphere, mean and standard deviations of T1w/T2w maps were 1.99 ± 0.43 for SUB, 2.01 ± 0.68 for CA, and 1.69 ± 0.23 for DG.

**Supplementary Fig. S4.**
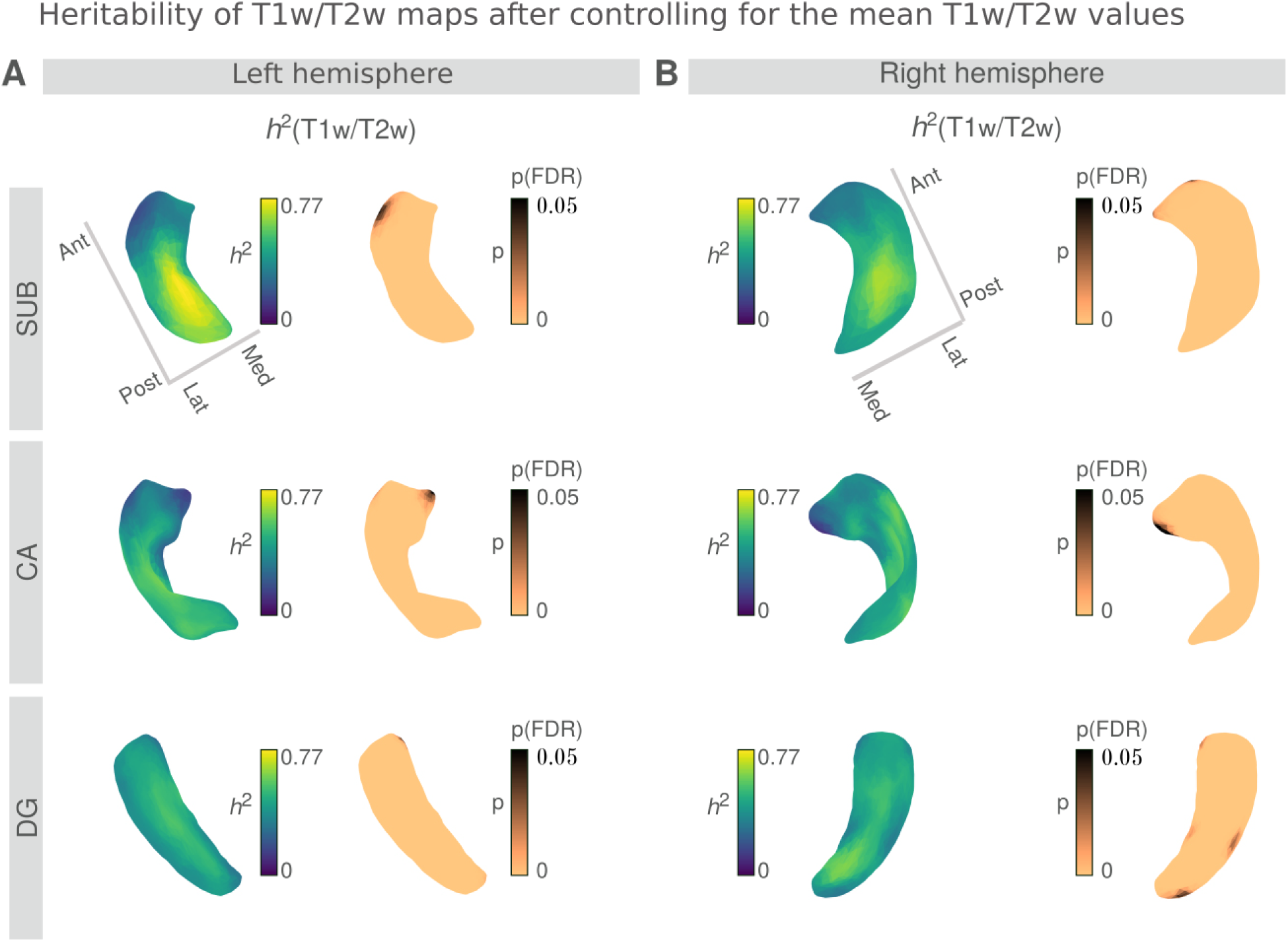
Heritability of the T1w/T2w intensity maps (*h*^2^(T1w/T2w)) by controlling for the mean T1w/T2w intensities for each subfield (SUB, CA, DG) and hemisphere (left and right). **A**. *h*^2^(T1w/T2w) patterns and their FDR-corrected significance levels (p(FDR)) did not change after controlling for the mean T1w/T2w intensities for the left hemisphere (copper color denotes pFDR < 0.05, black pFDR > 0.05). **B**. Similar results were observed for the right hemisphere.

**Supplementary Fig. S5.**
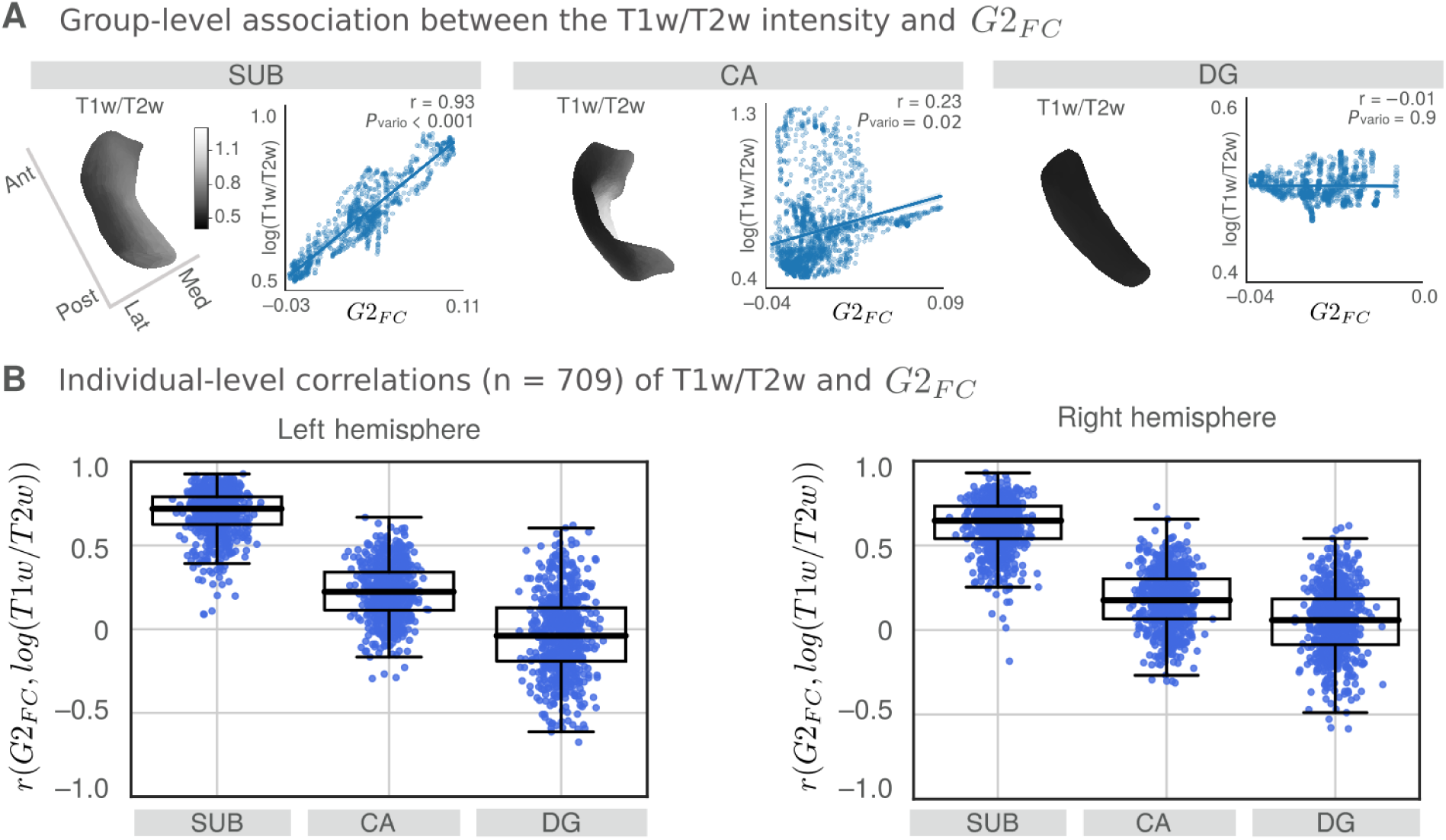
Spatial association between T1w/T2w intensity maps and the medial-lateral organisational axis represented by *G*2_*FC*_. **A**. Group-level association between the mean T1w/T2w profiles and the *G*2_*FC*_ for each subfield. The T1w/T2w profiles of subfields correlate strongly with the *G*2_*FC*_ for SUB (r = 0.93 and *p*_*vario*_< 0.001), however, not for the CA (r = 0.23 and *p*_*vario*_= 0.02) or DG (r = -0.01 and *p*_*vario*_= 0.9). **B**. Individual-level correlations (n = 709) between *G*2_*FC*_ maps and T1w/T2w intensity maps for each subfield (SUB, CA, DG) and hemisphere (left and right). For the left hemisphere, individual correlations (r(*G*2_*FC*_, log(T1w/T2w)) were significantly positive for the SUB (median 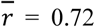, *p* < 0.005, one-tailed Wilcoxon signed-rank test) and CA (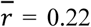, *p* < 0.005), however not for DG(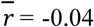, *p* < 0.01). Similar results were observed along the subfields in the right hemisphere 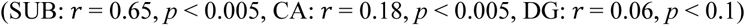.

## Notes

https://github.com/CNG-LAB/cngopen/tree/main/hippocampus

### Competing Interest Statement

The authors have declared no competing interest.

### Summary of Updates

Gradient alignment was erroneous, this has been corrected.

